# Ab-Ligity: Identifying sequence-dissimilar antibodies that bind to the same epitope

**DOI:** 10.1101/2020.03.24.004051

**Authors:** Wing Ki Wong, Sarah A. Robinson, Alexander Bujotzek, Guy Georges, Alan P. Lewis, Jiye Shi, James Snowden, Bruck Taddese, Charlotte M. Deane

## Abstract

Solving the structure of an antibody-antigen complex gives atomic level information of the interactions between an antibody and its antigen, but such structures are expensive and hard to obtain. Alternative experimental sources include epitope mapping and binning experiments which can be used as a surrogate to identify key interacting residues. However, their resolution is usually not sufficient to identify if two antibodies have identical interactions. Computational approaches to this problem have so far been based on the premise that antibodies with similar sequences behave similarly. Such approaches will fail to identify sequence-distant antibodies that target the same epitope.

We present Ab-Ligity, a structure-based similarity measure tailored to antibody-antigen interfaces. Using predicted paratopes on model antibody structures, we assessed its ability to identify those antibodies that target highly similar epitopes. Most antibodies adopting similar binding modes can be identified from sequence similarity alone, using methods such as clonotyping. In the challenging subset of antibodies whose sequences differ significantly, Ab-Ligity is still able to predict antibodies that would bind to highly similar epitopes (precision of 0.95 and recall of 0.69). We compared Ab-Ligity’s performance to an existing tool for comparing general protein interfaces, InterComp, and showed improved performance on antibody cases alongside a significant speed-up. These results suggest that Ab-Ligity will allow the identification of diverse (sequence-dissimilar) antibodies that bind to the same epitopes from large datasets such as immune repertoires. The tool is available at http://opig.stats.ox.ac.uk/resources.

## 1 Introduction

Antibodies are immune proteins that have high specificity and affinity against their target antigens. Their target specificity is determined by the intermolecular interactions at the antibody-antigen interface. The types of interactions used in antibody-antigen binding are known to be distinct from those observed in general protein-protein interactions [1].

The highest resolution method for studying antibody-antigen binding configurations is co-crystal complex structures. These give atomic level information but are expensive and difficult to obtain [2]. Experimental mapping is often used as a surrogate as it is able to identify the binding regions of the antigen (“epitopes”) and antibody (“paratopes”; [3]). Competition assays exploit the cross-blocking effect of antibodies that displace one another if they bind to similar or neighbouring epitopes [4, 5]. This method gives a coarse representation of which binders may share similar target sites, as minimal epitope overlap can be sufficient for a pair of antibodies to compete with each other [5]. A more refined approach is hydrogen deuterium exchange (HDX). HDX assesses the solvent accessibility of the bound and unbound forms of the partner proteins, and highlights regions with the maximum changes upon binding (*e.g.* [6, 7]). The resolution is typically up to the range of peptides in the immediate proximity of the binding site. To achieve residue-level resolution, point mutations of the interacting proteins can be used to indicate key binding residues. Mutagenesis studies measure the binding kinetics upon mutation of specific residues, but structural integrity may be compromised by the mutations, leading to spurious results [3]. All three of these experimental techniques provide an approximation of the binding regions, but are usually unable to provide a fine mapping of exact epitopes and paratopes.

Computational techniques have also been developed to identify antibodies that bind in similar ways. These have generally been exploited on large immunoglobulin sequencing datasets (*e.g.* [8]). Many of these techniques require a large number of known binders to a given epitope, to be able to identify further binders [9]. The informatics approaches employed to analyse datasets, when none or only a few binders to a given epitope are known, are mainly dependent on sequence similarity. This is based on the concept of “clonotype” analysis [8], which considers the genotype and the sequence identity of CDRH3 (the third complementarity determining region on the heavy chain) [8, 10, 11]. Clonotyping exploits the evolutionary origins of antibodies, using both the concept that antibodies from the same clone and lineage tend to bind similarly [12], and that the CDRH3 region, the most sequence-variable region, is often responsible for much of the binding [13]. Whilst clonotype analysis can identify antibodies with similar binding modes, there are many cases where binding remains the same even when the CDRH3 sequences and/or the genotypes are different (*e.g.* in anti-lysozyme antibodies; [14]).

To capture potential chemical interactions and the antibody-antigen binding configurations, macromolecular docking has been used [15]. However, this method is slow and not scalable to the large datasets of antibody sequences available in the early discovery stage [16].

In small molecule discovery, comparing the spatial arrangements of pharmacophores (the atom features involved in interactions) has been proposed as an alternative to docking when searching for similar binders. Pharmacophoric points are encoded using geometric hashing algorithms (*e.g.* [17–19]). This approach is much faster than docking whilst giving comparable results [19]. Some of the descriptors for ligand-binding pockets have been adapted for protein-protein interactions. The I2ISiteEngine uses atom type triangulation to describe protein-protein interfaces and is able to successfully identify similar interfaces across different protein families in a number of case studies [20]. However, the all-atom description means it is far slower than its counterpart developed for the comparison of small molecules [17]. Recently, a number of tools have been proposed for general protein-protein interface comparisons. MaSIF compares molecular surfaces of protein-protein interfaces using geometric deep learning [21]. Whilst this technique showed promise using bound structures to search for similar binders, the performance did not hold on a more challenging, unbound set. MaSIF’s high sensitivity to the precise location and chemical properties of the atoms suggests that it would depend heavily on model accuracy and would be unsuitable for use on modelled structures. InterComp has also been proposed for general protein-protein interface comparison. It aligns residues on the surfaces for comparison and scores their similarity using a combination of the C*α* atom distances and their BLOSUM substitution score [22]. The approach successfully identified similar surface patches but the algorithm is computationally intensive. Both of these methods were only applied on solved crystal structures and on general-protein-protein interfaces, not on models and more specifically, not on models of antibodies. In the context of finding antibodies in large sequencing datasets that target the same epitope, a tool needs to be fast, applicable on antibody-antigen interfaces, and able to cope with the predicted model structures of antibodies.

In this manuscript, we describe Ab-Ligity, an antibody version of the small molecule method Ligity [19]. Ab-Ligity uses residue points tokenized by their physicochemical properties to allow rapid operation. To simulate a real-life scenario, we applied Ab-Ligity to antibody models and predicted paratopes, and showed that it can accurately predict antibodies that share similar target epitopes. As the majority of the similar binders shared similar sequence composition and lengths, and can be identified by sequence-based metrics, we also considered the more challenging cases that could not be identified by such sequence-based metrics. In these more challenging cases of sequence-dissimilar antibodies, Ab-Ligity still accurately predicts antibodies that have similar binding modes. Ab-Ligity also performed better than InterComp, in a fraction of the time. Finally we describe two case studies where Ab-Ligity predicts sequence-dissimilar and length-mismatched antibodies that bind to highly similar epitopes.

## 2 Results

Ab-Ligity compares two antibody paratopes by tokenized residues and distance hashes. It is designed to work on antibody models with predicted paratopes. In this manuscript, we used Parapred [23] for paratope prediction (for full details, see Methods).

To test the power of Ab-Ligity to identify antibodies that bind to the same epitope, we simulated a real-life application of using antibody models and predicted paratopes. We built models of 920 unique protein-binding antibodies using ABodyBuilder [24], and predicted their paratope residues with Parapred [23]. Using these models and predicted paratopes we tested if Ab-Ligity was able to identify antibodies that bind to the same epitopes (as defined by the antibody-antigen crystal complexes; see Methods). We call this set the “full set”. Since the majority of the similar binders have the same CDRH3 length and highly similar CDRH3 sequences, we also tested Ab-Ligity performance when we removed these “easy” comparisons (CDRH3 identity >0.8). We call this set the CDRH3≤0.8 set. The number of comparisons for each of these tests are summarized in Table 1.

**Table 1:**
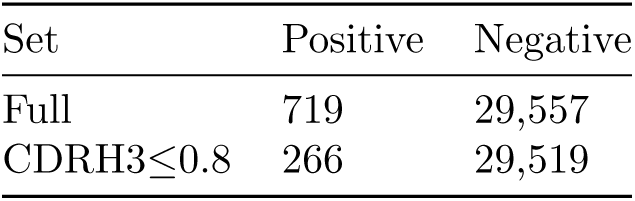
Number of positive and negative comparisons in the datasets, based on Ab-Ligity’s definition of similar epitopes.

| Set | Positive | Negative |
| --- | --- | --- |
| Full | 719 | 29,557 |
| CDRH3≤0.8 | 266 | 29,519 |

### 2.1 Selecting similarity thresholds

#### 2.1.1 Definition of similar epitopes captured in crystal structures

In order to identify a standardised definition of when two epitopes are the same, we ran a grid search on the corresponding “crystal paratope” and “crystal epitope” similarity scores as calculated by Ab-Ligity, maximising the Matthews correlation coefficient (MCC; see Methods and Supplementary Materials). This procedure suggested an epitope similarity threshold of 0.1 was appropriate and visual inspections of examples confirmed this. Epitope pairs above this threshold are considered positive (Table 1).

#### 2.1.2 Selecting a similarity threshold for predicted paratopes in antibody models

In a real-life scenario where antibody models and predicted paratopes would be used, we need to establish a model paratope similarity threshold that can recapitulate the definition of similar epitopes as defined by the crystal structures. Similar to the earlier strategy, we ran a grid search on a range of “model paratope” similarity scores, and found a threshold of 0.1 that gives the best MCC (0.90, see Supplementary Materials). Above the model paratope similarity of 0.1, paratopes are predicted to bind to a common epitope.

### 2.2 Using Ab-Ligity to predict antibodies that bind to highly similar epitopes

We assessed the performance of Ab-Ligity on two datasets of different difficulties: the “easy” full set and the “hard” CDRH3≤0.8 set. Sequence-based methods can identify relatively accurately CDRH3 sequence-similar antibodies that will bind to the same epitope [25]. In our dataset, the majority of the antibody pairs that target highly similar epitopes have similar CDRH3 sequences. Table 1 shows that 63.0% (453/719) of the positive pairs that bind to highly similar epitopes have CDRH3 sequence identities >0.8. This set represents “easy” cases, and Ab-Ligity is able to stratify between similar and dissimilar binders with good accuracy on the “easy” full set (precision of 0.95 and recall of 0.85; Table 2).

**Table 2:** Precision and recall on the full and CDRH3 ≤0.8 sets, using the selected paratope similarity thresholds for Ab-Ligity (0.1) and InterComp (0.6), based on Ab-Ligity’s definition of similar epitopes.

| Method | Ab-Ligity |  | InterComp |  |
| --- | --- | --- | --- | --- |
| Set | Precision | Recall | Precision | Recall |
| Full | 0.95 | 0.85 | 0.92 | 0.77 |
| CDRH3≤0.8 | 0.95 | 0.69 | 0.86 | 0.59 |

To assess Ab-Ligity’s performance on a more challenging set, we removed the “easy” comparisons. The remaining 37.0% (266/719; see Table 1) of the pairs that bind to similar epitopes have their CDRH3 sequence identities ≤0.8, and would not be identified by sequence-based methods such as clonotype. On this more challenging subset of cases, Ab-Ligity’s precision remains at 0.95 and its recall drops slightly (0.69; Table 2). These results demonstrate that Ab-Ligity is able to accurately identify sequence-dissimilar antibodies that have similar binding modes.

### 2.3 Sensitivity analyses

We assessed the sensitivity of Ab-Ligity’s performance in three situations: varying the distance bin size in hashing, changing the Parapred prediction threshold and applying Ab-Ligity to only the heavy (VH) or light chain (VL). The latter two factors change the paratope size. Using different Parapred thresholds affects the number of paratope residues predicted on each of the antibodies and potentially introduces noise if an inappropriate threshold is selected. We checked the performance on VH and VL alone as almost all current large sequencing datasets are unpaired [26]; if Ab-Ligity is able to accurately predict on a single chain, its potential application space is increased.

#### 2.3.1 Distance bin size

During the hashing procedure, Ab-Ligity discretizes the distances between residues (*i.e.* edge length of the triangles) into distance bins of 1Å. We observed that tightening the bin width to 0.5 Å marginally reduces the classification performance with a precision and recall of 0.93 and 0.85 in the full set and a similar change on the CDRH3≤0.8 set (see Supplementary Materials). Increasing the bin size harms performance, potentially as it over-smooths residue distances (see Supplementary Materials).

#### 2.3.2 Parapred predictions

Ab-Ligity uses predicted paratopes, so we examined how their accuracy influences performance. The Parapred prediction threshold of 0.67 from the original paper was selected such that the size of the predicted paratopes replicates that of actual paratopes. At this threshold, the precision and recall of Parapred were found to be 0.67 and 0.73 (see Supplementary Materials). We increased the threshold to 0.80 (increasing precision and reducing the number of residues predicted; see Supplementary Materials) to investigate the effect on Ab-Ligity. We saw little effect on the performance of Ab-Ligity: the precision and recall were 0.95 and 0.85 for the original Parapred threshold (0.67), compared to 0.96 and 0.80 in the increased Parapred thresholds (see Supplementary Materials).

We then reduced the Parapred threshold, increasing the paratope size but lowering the precision of paratope prediction. This reduction in the Parapred threshold decreased Ab-Ligity performance (see Supplementary Materials). This drop in performance is probably due to an elevated level of noise (*i.e.* the inclusion of residues not involved in binding) in the Parapred predictions.

#### 2.3.3 Performance on heavy chains or light chains alone

Since most publicly available immune repertoire datasets are from bulk sequencing of unpaired antibody chains [26], we assessed the applicability of Ab-Ligity on unpaired heavy and light chains.

We built homology models for heavy chain or light chain separately and extracted the predicted paratopes on these models. In Table 3, we show that heavy chain paratopes alone can be used to accurately identify antibodies that bind to the same epitopes, with a precision of 0.88 and recall of 0.78. Using the light chain alone, Ab-Ligity can also identify antibodies that bind to similar epitopes but with lower precision (Table 3). We also evaluated using the heavy chain or light chain paratopes from paired antibody models and the results were almost identical (see Supplementary Materials).

**Table 3:** Precision and recall on the full set, using heavy chain and light chain only homology models and the predicted paratopes on the corresponding single domain.

| Method | Ab-Ligity |  | InterComp |  |
| --- | --- | --- | --- | --- |
| Domain | Precision | Recall | Precision | Recall |
| VH | 0.88 | 0.78 | 0.74 | 0.82 |
| VL | 0.67 | 0.89 | 0.17 | 0.94 |

### 2.4 Comparing Ab-Ligity to InterComp

We compared Ab-Ligity to an existing general protein-protein interface comparison tool, InterComp. Since the original manuscript of InterComp did not indicate a threshold for interfaces to be considered similar, we conducted the same evaluation as for Ab-Ligity. The InterComp epitope similarity threshold was selected at 0.7 by maximising the MCC between crystal paratope and crystal epitope similarities (see Supplementary Materials). Based on this definition of similar epitopes, 0.6 is the optimal cut-off for InterComp model paratope similarity.

In Table 2, we show the results of Ab-Ligity and InterComp using Ab-Ligity’s definition of similar crystal epitopes. In the Supplementary Materials, we give the corresponding results for the methods using InterComp’s definition of similar epitopes. Both Table 2 and the Supplementary Tables show that the two methods have comparable performance with Ab-Ligity showing improved performance on the more challenging CDRH3 ≤ 0.8 set, regardless of whether the performance is assessed using the definition of ground truth by Ab-Ligity or InterComp (*i.e.* which pairs of antibody-antigen structures were considered to have similar epitopes by the two methods).

We carried out the same sensitivity analyses on InterComp as described above for Ab-Ligity. In terms of Parapred thresholds and predictions on VH or VL chains alone, very similar effects were observed (see Supplementary Materials).

As well as outperforming InterComp, Ab-Ligity is also significantly faster. We measured the algorithm run-time on a single 3.40GHz i7-6700 CPU core. On the full set of 920×920 pairwise comparisons, Ab-Ligity takes 0.5 CPU-minute to generate the 920 hash tables for all paratopes, and 18.5 minutes for an optimized all-against-all similarity calculation. InterComp does not require pre-processing but the all-against-all query takes 65.5 minutes. It is now possible to model large portions of next generation sequencing datasets [27] and Ab-Ligity would allow rapid comparison of binding sites in these datasets. For example, Ab-Ligity would take one day to process on 150 cores a next generation dataset of 100,000 antibodies, compared to five days for InterComp.

### 2.5 Anti-lysozyme antibodies with dissimilar CDRH3 sequences against highly similar epitopes

To show the power of Ab-Ligity to predict similar binding of antibodies with dissimilar CDRH3 sequences, we examined three anti-lysosyzme antibodies, HyHEL-26, HyHEL-10 L-Y50F mutant and HyHEL-63 (annotated as 1NDM_BA_C, 1J1O_HL_Y and 1NBY_BA_C respectively in Figure 1). These antibodies are known to all adopt the same binding mode against lysozyme (Figure 1A; [28]). Their CDRH3 loops are seven residues long, but differ by at least three residues between any pair, so would not be considered to share the same clonotype (Figure 1B). We have also included a counter-example of the 3C8 antibody (annotated as 6OKM_HL_R in Figure 1) against Tumor Necrosis Factor Receptor superfamily member 4 (OX4). This antibody also has seven-residue long CDRH3 sequence and shares a maximum sequence identity of 0.57 with HyHEL-26 (1NDM_BA_C).

**Figure 1:**
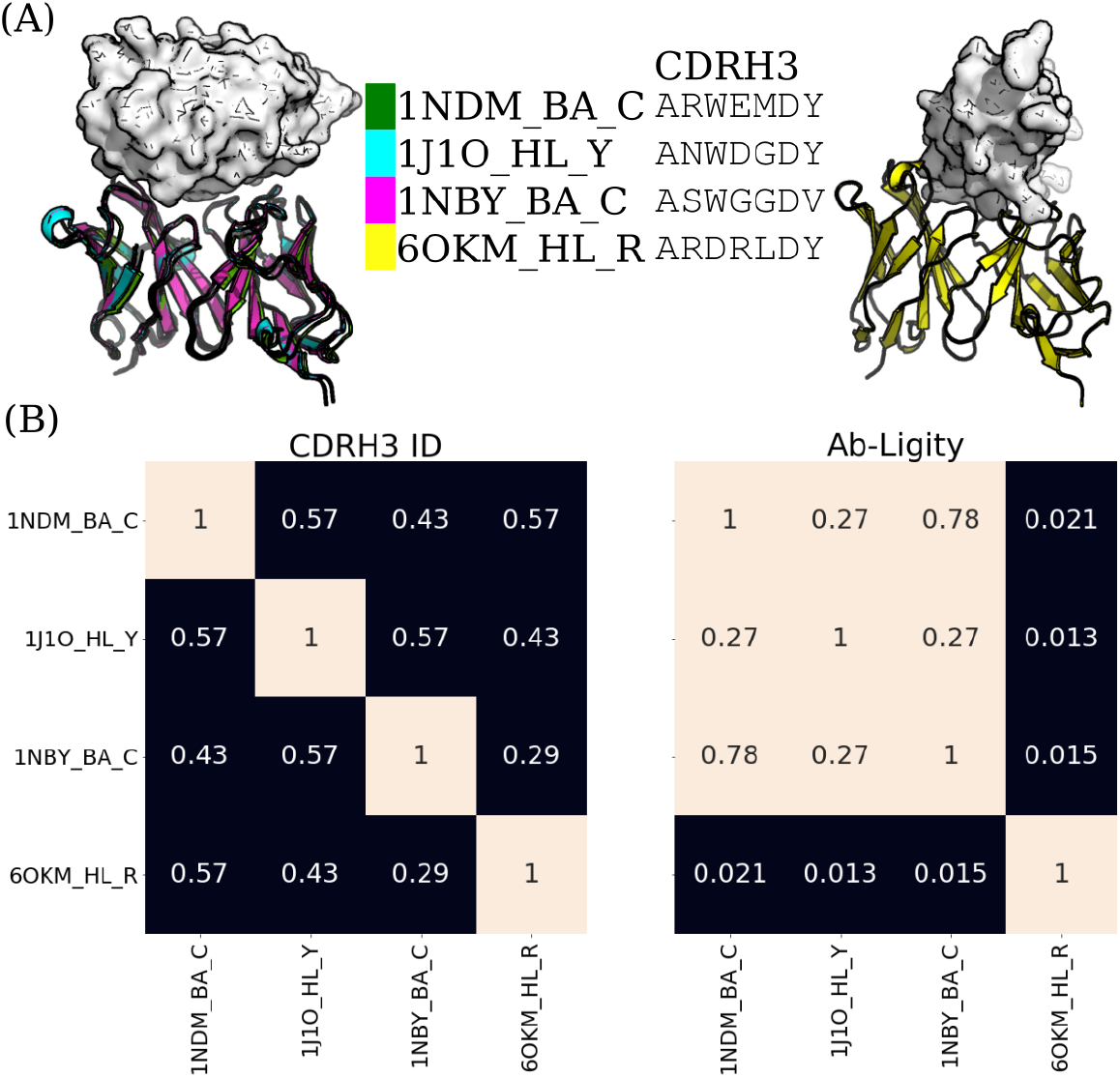
Analysis of anti-lysozyme antibodies with dissimilar CDRH3 sequences and highly similar epitopes. (A) Structural superposition of three anti-lysozyme and one anti-OX4 antibodies co-crystallized with their antigens (lysozyme or OX4) in white. The three anti-lysozyme antibodies were HyHEL-26 (1NDM_BA_C), HyHEL-10 L-Y50F mutant (1J1O_HL_Y) and HyHEL-63 (1NBY_BA_C); and the anti-OX4 antibody is 3C8 (6OKM_HL_R). The antigens from the three anti-lysozyme antibody crystal structures are aligned. The legend shows the colors of the antibodies with their PDB codes followed by the heavy-light chain and antigen chain identifiers, separated by (‘_’). The CDRH3 sequences are displayed next to the respective antibody identifiers. (B) Heatmaps of CDRH3 sequence identity and Ab-Ligity paratope similarity. The row and column labels correspond to the structures shown in (A). Ab-Ligity paratope similarity is calculated on the antibody model and predicted paratope as outlined in the manuscript. Pairs of antibodies with CDRH3 identity of >0.80 would have been considered similar by sequence-based metric. For Ab-Ligity, a similarity score of >0.1 suggests that the antibodies bind to highly similar epitopes.

We used Ab-Ligity to calculate the similarity of their binding using models of predicted paratopes. As outlined above, an Ab-Ligity score of above 0.1 indicates that the antibodies in comparison have similar binding sites. The pairwise similarity scores of all three anti-lysozyme antibodies are all above 0.1, indicating that Ab-Ligity would classify them as targeting the same epitope. This classification is consistent with the observation in the co-crystal complexes (Figure 1A). Conversely, the Ab-Ligity scores of these three anti-lysozyme antibodies with the anti-OX4 antibody were all below 0.1. Reflected in the crystal structures shown in Figure 1A, the anti-OX4 antibody clearly binds to a different antigen and epitope to the other three antibodies.

### 2.6 CDRH3 sequences with different lengths engage the same epitope in HIV core gp120

Ab-Ligity may also be useful in cases where antibodies have different CDRH3 lengths but similar binding modes. Sequence-based metrics are not applied to such cases [8]. One example of mis-matched CDRH3 lengths binding the same epitope are two HIV-1 neutralizing antibodies, VRC01 (PDB code and chain ID 4LSS_HL) and its variant, VRC07 (4OLU_HL). They target the gp120 core with highly similar binding modes (Figure 2). Their heavy and light chain germlines are the same but their CDRH3 sequences differ in length and composition. Aligning by the IMGT-numbered positions [29], only 12 out of the 18 residues are aligned and identical. These two antibodies are known to engage a similar epitope [30] and Ab-Ligity based on models of these antibodies and predicted paratopes correctly predicts this (Ab-Ligity score of 0.24; ≥0.1 Ab-Ligity similarity threshold).

**Figure 2:**
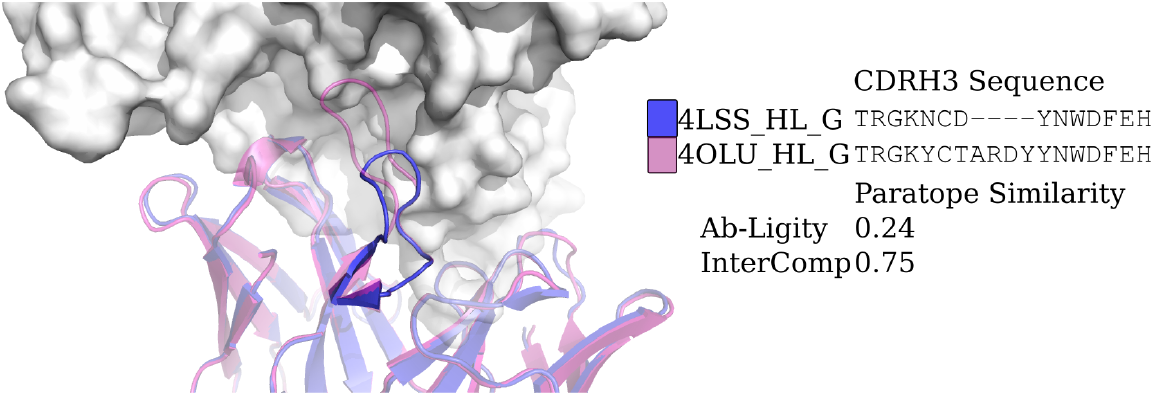
Analysis of two anti-gp120 antibodies. VRC01 antibody (PDB code and chain IDs: 4LSS_HL_G) is colored in blue; VRC07 (PDB code and chain IDs: 4OLU_HL_G) is shown in pink. The gp120 core antigen is displayed as white surfaces and the superimposed antibodies are in cartoons. The CDRH3 loops are in solid shades of the cartoon representation. The PDB code, heavy and light chain ID, and antigen chain ID are separated by (‘_’) and listed in the legend with the corresponding CDRH3 sequences. CDRH3 sequences are shown by aligning their IMGT positions and ‘-’ indicates a gap in the alignment according to the IMGT numbering scheme [29]. The Ab-Ligity paratope similarity score for the pair is listed.

## 3 Discussion

We present Ab-Ligity, an antibody-protein binding site structural similarity metric that can identify sequence-dissimilar antibodies that engage the same epitope. Our results show that Ab-Ligity is able to identify antibodies that bind to the same epitopes using model structures and predicted paratopes, even for pairs of sequence-dissimilar antibodies. We evaluated the robustness of Ab-Ligity to the distance hashing, its dependence on the accuracy of paratope prediction by Parapred, and its applicability on unpaired antibody sequencing datasets. We also compared Ab-Ligity to InterComp, an existing protein surface similarity metric and found improved performance for harder cases with dissimilar sequences, and for the application on heavy chains or light chains only paratopes, and a far faster run-time. We further show that Ab-Ligity can identify antibodies that bind to the same epitope with dissimilar CDRH3 lengths, beyond the constraints of most sequence-based metrics.

Binding site comparison with tools, such as Ab-Ligity, opens up an alternative way to search for binders with similar binding modes. Typically in antibody discovery, multiple diverse hits are desired to avoid developability issues in the downstream optimisation process [31]. Ab-Ligity has the ability to predict if antibodies share similar target epitopes, without being sequence-similar. This ability coupled with the fast run-time of Ab-Ligity makes it suitable for searching through large datasets of antibody sequences to find sets of sequence-diverse binders to the same epitope.

## 4 Methods

### 4.1 Antibody-antigen co-crystal datasets

We selected all paired antibody-antigen complexes from SAbDab [2] as of 27th January, 2020, that were solved by X-ray crystallography, had no missing residues in all six CDRs (using the North definition; [13]), and were co-crystallized with a protein antigen of more than 50 residues. To avoid redundancy, we retain only one copy of each antibody. In the case of multiple copies, the complex with the best resolution is selected. Nine hundred and twenty antibody-antigen complexes were identified.

We defined epitopes as residues on the antigen with any atoms within 4.5Å of its cognate antibody. We extracted these residues using Biopython [32].

### 4.2 Antibody modelling and paratope prediction

We numbered antibody sequences using the IMGT scheme [29] and defined the CDR regions by the North scheme [13], with ANARCI [33]. We then modelled the full set using ABodyBuilder [24], barring it from using sequence-identical structures as templates. The ABodyBuilder template library was built using all structures available on the 27th January, 2020, in SAbDab [2].

We used Parapred to predict the paratope residues [23]. Parapred gives a score to each residue in the CDRs, and two residues before and after, to indicate how likely it is to participate in binding. As suggested in the original paper, we selected residues with a score of ≥0.67 as the predicted paratope residues on the models (see Supplementary Materials for Parapred prediction performance). The co-ordinates of these predicted paratope residues were obtained from the corresponding model. Models and predicted paratopes were used throughout the manuscript for calculating the structural similarity between antibodies.

### 4.3 Ab-Ligity calculations

The workflow of Ab-Ligity is illustrated in Figure 3. The binding residues are tokenized according to Table 4. This tokenisation scheme was chosen as it is simple and intuitive, and gave similar results to several other more complex choices. For each binding site residue, a point is placed representing the residue group on the C-*α* atom of the residue. These points collectively represent a paratope on an antibody or an epitope on an antigen.

**Table 4:** Residue groupings for tokenisation.

| Group | Residues |
| --- | --- |
| Aliphatic | Glycine (G), Alanine (A), Valine (V), Leucine (L), Isoleucine (I), Proline (P) |
| Hydroxyl | Serine (S), Threonine (T) |
| Sulphur | Cysteine (C), Methionine (M) |
| Aromatic | Phenylalanine (F), Tyrosine (Y), Tryptophan (W) |
| Acidic | Aspartic acid (D), Glutamic acid (E) |
| Amine | Asparagine (N), Glutamine (Q) |
| Basic | Histidine (H), Lysine (K), Arginine (R) |

**Figure 3:**
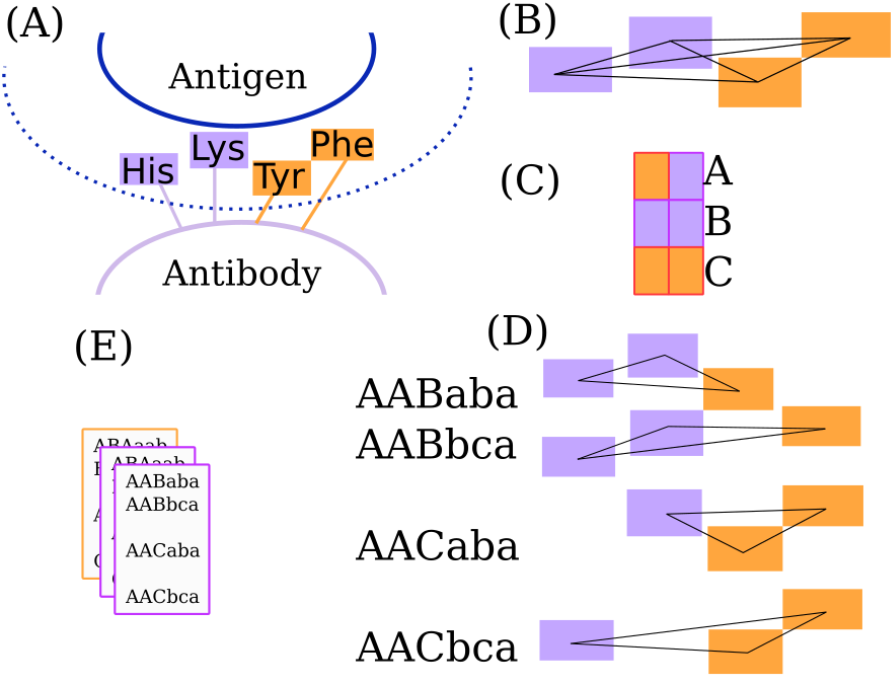
The Ab-Ligity workflow. (A) Binding site residues within 4.5Å of the binding partner are tokenized as stated in Table 4. (B) All distances between tokenized points are calculated and hashed into 1.0Å-wide distance bins for both the paratopes and epitopes. (C) Each pair of tokens is given a unique hash code. (D) A six-character hash code is generated for each triplet. (E) The hashes of a binding site are stored in a frequency table.

Ab-Ligity uses the hashing function outlined in [19]. In short, it considers all combinations of triplets formed from a set of tokenized residues in a binding site. In a given triplet, each edge is represented by its vertices’ tokens and its length. Each combination of tokens has a unique hash code. Edge lengths are discretized into bins of 1.0Å in both paratopes and epitopes to reduce computational complexity. The final hash code for a given triplet is determined by the three vertex hash codes sorted alphabetically, followed by the hash codes of the three corresponding length bins. The resulting hash table for the target binding site stores the frequency of these hash codes from all triplets.

Binding site similarity is calculated by the Tversky index of the pairs of hash tables:

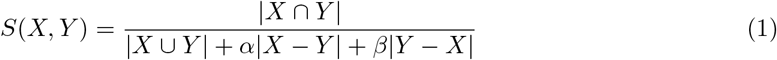

where *X* and *Y* are the two hash tables, *|X ∩ Y|* and *|X ∪ Y|* are the intersection and union, and *|X − Y|* and *|Y − X|* are the differences between the two tables respectively. For this study we used *α* = *β* = 0.5.

### 4.4 Performance evaluation settings

We calculated the Ab-Ligity paratope similarity based on antibody models and their predicted paratopes. The majority of the antibodies that engage the same epitope have highly similar CDRH3 sequences and would be predicted to bind in the same manner by sequence comparison (such as clonotype). To test if Ab-Ligity makes accurate predictions for those cases without similar CDRH3s, we assessed the performance on both the “full set”, and a subset where the CDRH3 identity is ≤ 0.8 (“CDRH3≤0.8 set”).

The number of positive and negative cases for these two sets are listed in Table 1. The performance at the selected thresholds is reported as precision and recall (see Supplementary Materials).

### 4.5 Selecting an epitope similarity threshold

For each pair of antibody-antigen complexes, we first calculated the epitope similarity based on the crystal structures of the antibody-antigen complexes. We selected the epitope similarity threshold by evaluating pairs of proposed crystal paratope and crystal epitope similarity thresholds (see Supplementary Materials) and chose the epitope similarity score that gives the best classification performance, *i.e.* the highest Matthews correlation coefficient (MCC; see Supplementary Materials). This becomes the ground truth for evaluation. For Ab-Ligity, we selected the crystal epitope similarity threshold at 0.1. (See Supplementary Materials for the corresponding comparisons using InterComp’s definition of similar epitopes.) Manual inspection of example cases indicated that pairs of epitopes above this threshold were highly similar. This threshold is used throughout the manuscript to give the binary classification of similar and dissimilar epitopes.

### 4.6 Benchmark

We compared our prediction performance and computational time with InterComp, a surface comparison tool for protein-protein interfaces [22]. (See Supplementary Materials for the corresponding analysis using InterComp’s definition.) The algorithm run-time was measured on a single 3.40GHz i7-6700 CPU core.

## Acknowledgement(s)

The authors thank Dr. Jean-Paul Ebejer for providing the original implementation of Ligity, Matthew Raybould for useful discussion of the residue tokenisation strategy, and members at Roche and GlaxoSmithKline for helpful discussion.

## References

[1] Wang M, Zhu D, Zhu J, et al. Local and global anatomy of antibody-protein antigen recognition. Journal of Molecular Recognition. 2018;31(5):e2693.

[2] Dunbar J, Krawczyk K, Leem J, et al. SAbDab: the structural antibody database. Nucleic Acids Research. 2013;42(D1):D1140–D1146.

[3] Abbott WM, Damschroder MM, Lowe DC. Current approaches to fine mapping of antigen– antibody interactions. Immunology. 2014;142(4):526–535.

[4] Kwak JW, Yoon CS. A convenient method for epitope competition analysis of two monoclonal antibodies for their antigen binding. Journal of Immunological Methods. 1996;191(1):49–54.

[5] Abdiche YN, Yeung AY, Ni I, et al. Antibodies targeting closely adjacent or minimally overlapping epitopes can displace one another. PloS One. 2017;12(1):e0169535.

[6] Zhang Q, Yang J, Bautista J, et al. Epitope mapping by HDX-MS elucidates the surface coverage of antigens associated with high blocking efficiency of antibodies to birch pollen allergen. Analytical Chemistry. 2018;90(19):11315–11323.

[7] Puchades C, Kűkrer B, Diefenbach O, et al. Epitope mapping of diverse influenza Hemagglutinin drug candidates using HDX-MS. Scientific Reports. 2019;9(1):4735.

[8] Galson JD, Trück J, Fowler A, et al. In-depth assessment of within-individual and interindividual variation in the b cell receptor repertoire. Frontiers in Immunology. 2015;6:531.

[9] Mason DM, Friedensohn S, Weber CR, et al. Deep learning enables therapeutic antibody optimization in mammalian cells by deciphering high-dimensional protein sequence space. bioRxiv. 2019;:617860.

[10] Trück J, Ramasamy MN, Galson JD, et al. Identification of antigen-specific B cell receptor sequences using public repertoire analysis. The Journal of Immunology. 2015;194(1):252–261.

[11] Soto C, Bombardi RG, Branchizio A, et al. High frequency of shared clonotypes in human B cell receptor repertoires. Nature. 2019;566(7744):398.

[12] Hsiao YC, Chen YJJ, Goldstein LD, et al. Restricted epitope specificity determined by variable region germline segment pairing in rodent antibody repertoires. mAbs. 2020;12(1):1722541.

[13] North B, Lehmann A, Dunbrack Jr RL. A new clustering of antibody CDR loop conformations. Journal of Molecular Biology. 2011;406(2):228–256.

[14] Pons J, Stratton JR, Kirsch JF. How do two unrelated antibodies, hyhel-10 and f9. 13.7, recognize the same epitope of hen egg-white lysozyme? Protein Science. 2002;11(10):2308–2315.

[15] Lensink MF, Nadzirin N, Velankar S, et al. Modeling protein-protein, protein-peptide and protein-oligosaccharide complexes: CAPRI 7th edition. Proteins: Structure, Function, and Bioinformatics. 2019;.

[16] Raybould MIJ, Wong WK, Deane CM. Antibody-antigen complex modelling in the era of immunoglobulin repertoire sequencing. Molecular Systems Design & Engineering. 2019;4(4):679–688.

[17] Shulman-Peleg A, Nussinov R, Wolfson HJ. SiteEngines: recognition and comparison of binding sites and protein–protein interfaces. Nucleic Acids Research. 2005;33(Suppl 2):W337–W341.

[18] Wood DJ, Vlieg Jd, Wagener M, et al. Pharmacophore fingerprint-based approach to binding site subpocket similarity and its application to bioisostere replacement. Journal of Chemical Information and Modeling. 2012;52(8):2031–2043.

[19] Ebejer JP, Finn PW, Wong WK, et al. Ligity: A non-superpositional, knowledge-based approach to virtual screening. Journal of Chemical Information and Modeling. 2019;59(6):2600–2616.

[20] Shulman-Peleg A, Mintz S, Nussinov R, et al. Protein-protein interfaces: Recognition of similar spatial and chemical organizations. In: International Workshop on Algorithms in Bioinformatics; Springer; 2004. p. 194–205.

[21] Gainza P, Sverrisson F, Monti F, et al. Deciphering interaction fingerprints from protein molecular surfaces using geometric deep learning. Nature Methods. 2020;17(2):184–192.

[22] Mirabello C, Wallner B. Topology independent structural matching discovers novel templates for protein interfaces. Bioinformatics. 2018;34(17):i787–i794.

[23] Liberis E, Veličcković P, Sormanni P, et al. Parapred: antibody paratope prediction using convolutional and recurrent neural networks. Bioinformatics. 2018;34(17):2944–2950.

[24] Leem J, Dunbar J, Georges G, et al. ABodyBuilder: Automated antibody structure prediction with data–driven accuracy estimation. mAbs. 2016;8(7):1259–1268.

[25] Scheid JF, Mouquet H, Ueberheide B, et al. Sequence and structural convergence of broad and potent hiv antibodies that mimic cd4 binding. Science. 2011;333(6049):1633–1637.

[26] Kovaltsuk A, Leem J, Kelm S, et al. Observed antibody space: A resource for data mining nextgeneration sequencing of antibody repertoires. The Journal of Immunology. 2018;201(8):2502–2509.

[27] Raybould MI, Marks C, Kovaltsuk A, et al. Evidence of antibody repertoire functional convergence through public baseline and shared response structures. BioRxiv. 2020;Available from: https://www.biorxiv.org/content/10.1101/2020.03.17.993444v1.

[28] Mohan S, Sinha N, Smith-Gill SJ. Modeling the binding sites of anti-hen egg white lysozyme antibodies hyhel-8 and hyhel-26: an insight into the molecular basis of antibody cross-reactivity and specificity. Biophysical journal. 2003;85(5):3221–3236.

[29] Lefranc MP, Giudicelli V, Duroux P, et al. IMGT®, the international ImMunoGeneTics information system® 25 years on. Nucleic Acids Research. 2014;43(D1):D413–D422.

[30] Li Y, O’Dell S, Wilson R, et al. HIV-1 neutralizing antibodies display dual recognition of the primary and coreceptor binding sites and preferential binding to fully cleaved envelope glycoproteins. Journal of Virology. 2012;86(20):11231–11241. Available from: https://jvi.asm.org/content/86/20/11231.

[31] Raybould MIJ, Marks C, Krawczyk K, et al. Five computational developability guidelines for therapeutic antibody profiling. Proceedings of the National Academy of Sciences. 2019; 116(10):4025–4030.

[32] Cock PJA, Antao T, Chang JT, et al. Biopython: freely available python tools for computational molecular biology and bioinformatics. Bioinformatics. 2009;25(11):1422–1423.

[33] Dunbar J, Deane CM. ANARCI: antigen receptor numbering and receptor classification. Bioinformatics. 2016;32(2):298–300.

