## Supplementary Materials for "Ab-Ligity: Identifying sequence-dissimilar antibodies that bind to the same epitope"

#### S1 Ab-Ligity pipeline

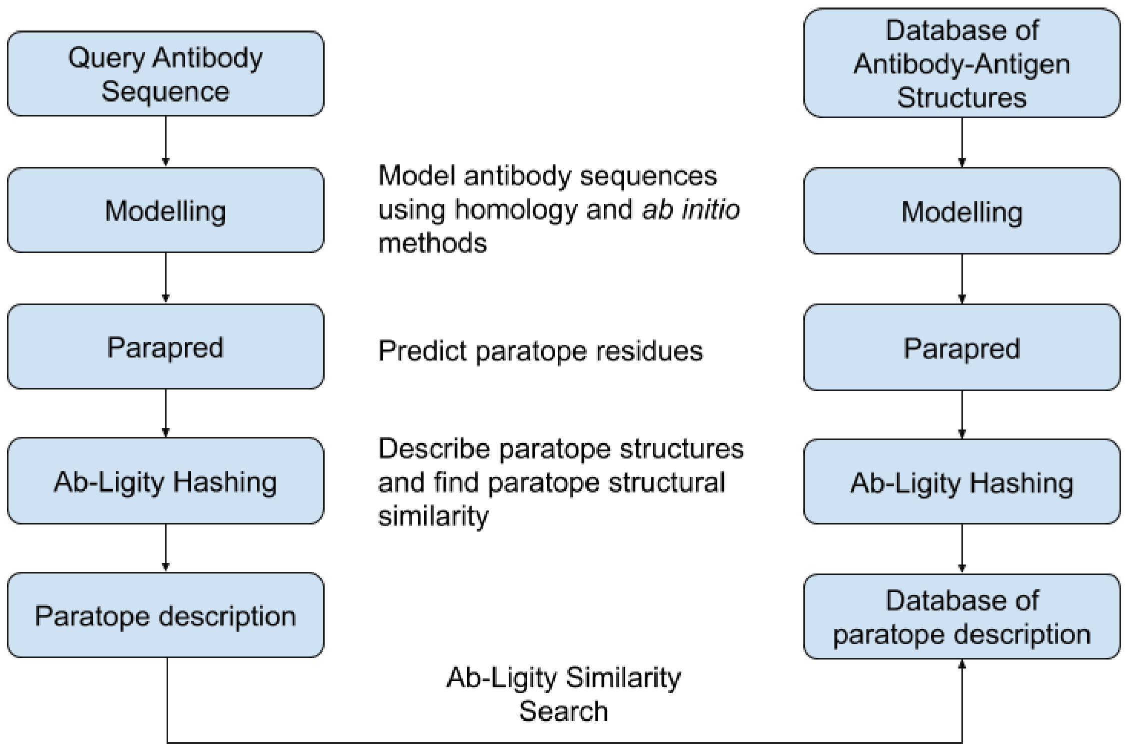


Figure S1. Ab-Ligity pipeline. A database of antibody-antigen structures is modelled, by ABodyBuilder in this manuscript (Leem *et al.*, 2016), has their paratopes predicted by Parapred (Liberis *et al.*, 2018), and hashed by the Ab-Ligity algorithm described in the Methods section of the manuscript. This yields a database of paratope description in the form of Ab-Ligity hash tables. For a query antibody sequence, it would undergo homology modelling, paratope prediction and Ab-Ligity hashing to produce a paratope description. This query paratope description is then used to query a database of paratope description to find the Ab-Ligity similarity scores against known paratopes.

### S2 Performance evaluation

#### S2.1 Selecting epitope similarity threshold

To select an epitope similarity threshold, we carried out an evaluation based on the paratopes and epitopes of the co-crystal complexes. Paratopes in the crystal structures (“crystal paratopes”) are defined as the antibody residues that have at least one atom within 4.5Å of the cognate antigen; likewise for the “crystal epitopes”. A set of hash tables and similarity scores were generated for the crystal paratopes and epitopes in the same way as for model paratopes (see manuscript).

We tested the classification performance by sweeping through pairs of crystal paratope and crystal epitope similarities in increments of 0.1, between 0 to 1. Consider a pair of antibodies with paratope similarity $S_p$ and corresponding epitope similarity of $S_e$ :

- True Positive (TP): $S_p \geq S_p{}^t$ and $S_e \geq S_e{}^t$
- True Negative (TN): $S_p < S_p{}^t$ and $S_e < S_e{}^t$
- False Positive (FP): $S_p \geq S_p{}^t$ and $S_e < S_e{}^t$
- False Negative (FN): $S_p < S_p{}^t$ and $S_e \geq S_e{}^t$

To select the optimal epitope similarity threshold that has the best classification performance, we used the following definition for Matthews correlation coefficient (MCC) to evaluate this performance:

$$MCC = \frac{TP \times TN - FP \times FN}{\sqrt{(TP+FP)(TP+FN)(TN+FP)(TN+FN)}}$$

The epitope similarity score with the highest MCC is selected: 0.1 Ab-Ligity score for Ab-Ligity’s definition of similar epitopes, and 0.7 InterComp score for that of InterComp.

Table S1. Performance of the selected thresholds based on crystal paratope and crystal epitope similarities defined by the same method.

| **Methods** | **Paratope Similarity** | **Epitope Similarity** | **MCC** | **Precision** | **Recall** |
|---|---|---|---|---|---|
| Ab-Ligity | 0.1 | 0.1 | 0.94 | 0.98 | 0.89 |
| InterComp | 0.7 | 0.7 | 0.88 | 0.90 | 0.86 |

#### S2.2 Selecting model paratope similarity threshold for a real-life application

In a common real-life scenario where antibody models and predicted paratopes were used, we need to establish a “model paratope” threshold that can recapitulate epitope similarity as defined in the crystal structures. As above, we used MCC to define the model paratope threshold that should be used with antibody models and predicted paratopes. We selected 0.1 for Ab-Ligity, based on Ab-Ligity’s definition of similar crystal epitopes (0.1). For InterComp, we selected 0.6 as the model paratope similarity threshold, using InterComp’s definition of similar crystal epitopes (0.7).

Table S2. Performance of the selected thresholds based on model paratope and crystal epitope similarities defined by the same method.

| **Methods** | **Paratope Similarity** | **Epitope Similarity** | **MCC** | **Precision** | **Recall** |
|---|---|---|---|---|---|
| Ab-Ligity | 0.1 | 0.1 | 0.90 | 0.95 | 0.85 |
| InterComp | 0.6 | 0.7 | 0.81 | 0.80 | 0.83 |

#### S2.3 Evaluating classification performance on datasets

To find out the classification performance of both Ab-Ligity and InterComp, we used the following definitions for precision and recall:

$$Precision = \frac{TP}{TP+FP}$$

$$Recall = \frac{TP}{TP+FN}$$

### S3 Parapred performance

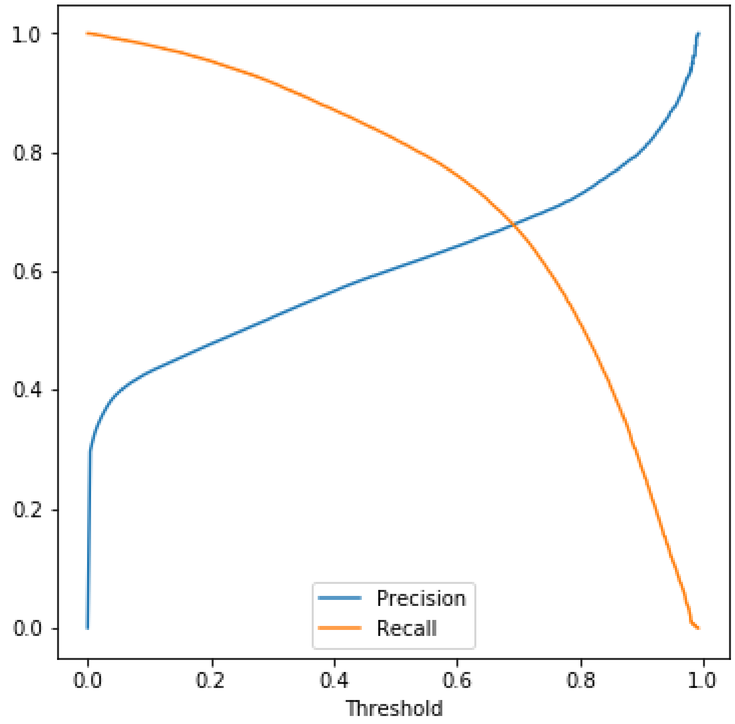


Figure S2. Performance of Parapred across the threshold. Precision and recall are defined as in S2.2.

In the manuscript, we used Parapred to predict the paratopes (Liberis *et al.*, 2018). Parapred gives a score to each residue in the CDRs, and two residues before and two after, to indicate how likely it will participate in binding. The precision and recall across different Parapred thresholds as calculated in the original paper (Liberis *et al.*, 2018) are presented in Figure S2.

In the original Parapred paper, they selected a threshold of 0.67 to balance between the actual and predicted paratope sizes. In the manuscript, we used this threshold to set the predicted paratopes for the Ab-Ligity and InterComp calculations. To test the sensitivity of these tools to the Parapred thresholds, we tested Parapred thresholds of 0.5 and 0.8 (see Section S6.2).

### S4 Performance on Ab-Ligity's definition

At our definition of epitope similarity (*i.e.* Ab-Ligity score of 0.1), we evaluated the performance using area under the precision-recall curve (AUPRC). At the selected paratope similarity thresholds, 0.1 for Ab-Ligity and 0.6 for InterComp, we showed the precision and recall.

Table S3. Performance in AUPRC on the core sets.

| **Set** | **Ab-Ligity** | **InterComp** |
|---|---|---|
| Full | 0.89 | 0.89 |
| CDRH3 ≤ 0.8 | 0.75 | 0.72 |

Table S4. Performance at the selected thresholds on the core sets.

| | **Ab-Ligity** | | **InterComp** | |
|---|---|---|---|---|
| **Set** | **Precision** | **Recall** | **Precision** | **Recall** |
| Full | 0.95 | 0.85 | 0.92 | 0.77 |
| CDRH3 ≤ 0.8 | 0.95 | 0.69 | 0.86 | 0.59 |

### S5 Performance on InterComp's definition

At InterComp's definition of epitope similarity (*i.e.* InterComp score of 0.7),we evaluated the performance using area under the precision-recall curve (AUPRC). At the selected paratope similarity thresholds, 0.1 for Ab-Ligity and 0.6 for InterComp, we showed the precision and recall.

Table S5. Number of positive and negative comparisons in the datasets, based on InterComp's definition of similar epitopes.

| **Set** | **Positive** | **Negative** |
|---|---:|---:|
| Full | 578 | 29,698 |
| CDRH3 ≤ 0.8 | 193 | 29,592 |

Table S6. Performance in AUPRC on the core sets.

| **Set** | **Ab-Ligity** | **InterComp** |
|---|---:|---:|
| Full | 0.83 | 0.81 |
| CDRH3 ≤ 0.8 | 0.73 | 0.69 |

Table S7. Performance at the selected thresholds on the core sets.

| | **Ab-Ligity** | | **InterComp** | |
|---|---:|---:|---:|---:|
| **Set** | **Precision** | **Recall** | **Precision** | **Recall** |
| Full | 0.83 | 0.92 | 0.80 | 0.83 |
| CDRH3 ≤ 0.8 | 0.81 | 0.82 | 0.73 | 0.69 |

### S6 Sensitivity analyses

#### S6.1 Distance bins

We tested the performance of Ab-Ligity when we used different bin sizes for the edge distance. For each distance bin size, we fixed the Ab-Ligity's definition of similar epitopes to a score of 0.1, and found the corresponding Ab-Ligity score with the highest Matthews' correlation coefficient at the respective thresholds. For the bin size of 0.5 Å, we kept the Ab-Ligity threshold for similar paratopes at 0.1, but we saw a slight increase in the Ab-Ligity threshold to 0.2 in the larger bin sizes.

Our current pipeline uses 1.0 Å for the distance bin hashing. In Table S8, we observed that the performance changes were negligible when decreasing the bin size to 0.5 Å or increasing to 1.5 Å. However, increasing the bin size to 2.0 Å harmed the precision, potentially due to over-smoothing.

Table S8. Performance of Ab-Ligity using different distance bin sizes on the two core sets, based on Ab-Ligity's definition of similar epitopes.

| **Distance bin size** | **0.5 Å** | | **1.0 Å (Original)** | | **1.5 Å** | | **2.0 Å** | |
|---|---|---|---|---|---|---|---|---|
| **Ab-Ligity threshold** | 0.1 | | 0.1 | | 0.2 | | 0.2 | |
| | Precision | Recall | Precision | Recall | Precision | Recall | Precision | Recall |
| **Full** | 0.93 | 0.85 | 0.95 | 0.85 | 0.98 | 0.74 | 0.82 | 0.56 |
| **CDRH3 ≤ 0.8** | 0.94 | 0.73 | 0.95 | 0.69 | 0.96 | 0.53 | 0.61 | 0.29 |

#### S6.2 Parapred thresholds

Since Ab-Ligity was developed to be used on predicted paratopes, we tested the effect of changing the Parapred threshold on the accuracies of Ab-Ligity and InterComp. The current Parapred threshold used in the manuscript is 0.67. We arbitrarily selected Parapred thresholds of 0.50 and 0.80 for this evaluation.

Reducing the Parapred threshold increased the number of residues in the CDR being predicted as the paratope, that is, the predicted paratopes became larger (Figure S3). This would also accentuate the noise by making more false positive predictions (Table S9). Under the threshold of 0.50, the performances of both Ab-Ligity and InterComp suffered (Table S10) because of the noise generated in the prediction.

On the contrary, increasing the Parapred threshold reduced the paratope sizes (Figure S3). Table S10 shows that Ab-Ligity and InterComp were only marginally insensitive to this surface size reduction.

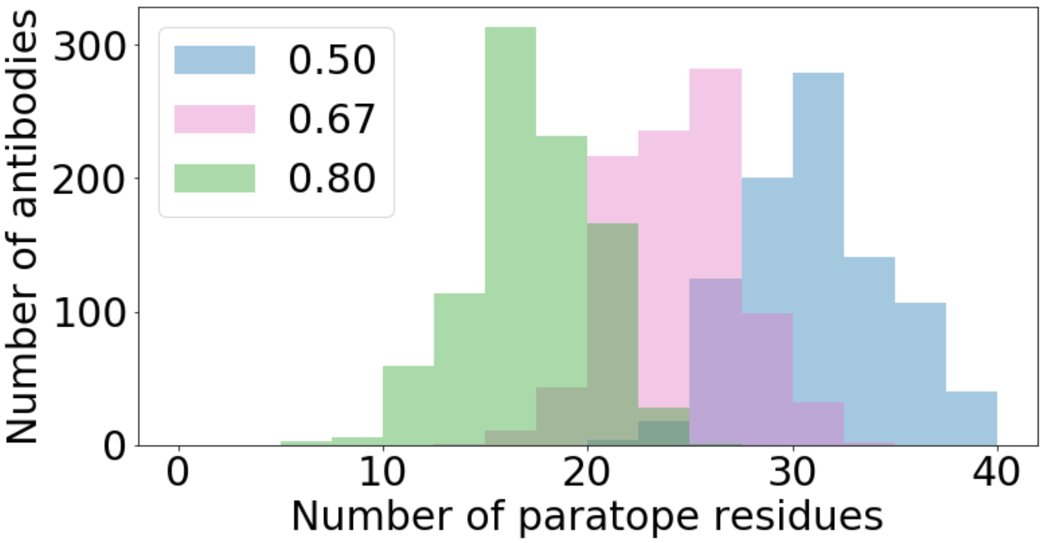


Figure S3. The size of the predicted paratopes in the non-redundant set, at different Parapred thresholds.

Table S9. Precision and recall of Parapred at selected thresholds.

| **Parapred Threshold** | **Parapred Performance** | |
|---|---|---|
| | **Precision** | **Recall** |
| 0.50 | 0.61 | 0.84 |
| 0.67 (Original) | 0.67 | 0.73 |
| 0.80 | 0.73 | 0.55 |

Table S10. Performance of Ab-Ligity and InterComp with predicted paratopes extracted at different Parapred thresholds, on the full set. The paratope similarity scores for Ab-Ligity and InterComp were 0.1 and 0.6 respectively; while the epitope similarity scores were 0.1 and 0.7 for Ab-Ligity and InterComp.

| | | **Epitope Definition** | | | |
|---|---|---|---|---|---|
| **Parapred Threshold** | | **Ab-Ligity** | | **InterComp** | |
| | **Paratope Definition** | **Precision** | **Recall** | **Precision** | **Recall** |
| **0.50** | **Ab-Ligity** | 0.73 | 0.25 | 0.66 | 0.28 |
| | **InterComp** | 0.87 | 0.23 | 0.79 | 0.25 |
| **0.67 (Original)** | **Ab-Ligity** | 0.95 | 0.85 | 0.83 | 0.92 |
| | **InterComp** | 0.92 | 0.77 | 0.80 | 0.84 |
| **0.80** | **Ab-Ligity** | 0.96 | 0.80 | 0.84 | 0.87 |
| | **InterComp** | 0.96 | 0.73 | 0.83 | 0.79 |

#### S6.3 Predictions on heavy chains or light chains only

The majority of the currently-available immune repertoire datasets contains unpaired VH and VL sequences (Kovaltsuk *et al.*, 2018). To assess the applicability of Ab-Ligity and InterComp these datasets, we tested the classification performances of Ab-Ligity and InterComp on heavy chains or light chains only.

We carried out two tests. First we followed the original process outlined in the manuscript to build paired homology models of antibodies using both the VH and VL sequences. We then extracted the predicted paratope residues in each of the VH or VL chain, for the construction of paratope surfaces by Ab-Ligity and InterComp. The second test involved building homology models using a single VH or VL sequence, and taking the same set of paratope predictions as in the first test. The distinction of the second test arises from the coordinates of the paratope residues: these coordinates could be different from the paired models as the 'companion' chain was not present when calculating the structural clashes in the homology modelling process.

The performance of Ab-Ligity and InterComp is shown in Table S11. The full crystal epitope similarity (*i.e.* using the crystal epitope extracted from the paired, crystal structures) was considered to be the ground truth. On the full antibody heavy chain/light chain-only paratopes set, we saw that the performance of both methods using VH paratopes retained, whereas that of VL paratopes dropped. VH paratope similarity alone appeared to be sufficient to identify similar epitopes, although paratopes on VL still held a portion of the key information to determine the overall paratope similarity. A similar trend was observed on the single domain model, heavy chain or light chain only paratope sets (Table S11). In the case of VH-/VL-only paratopes, Ab-Ligity consistently outperforms InterComp. Mirabello and Wallner (2018) reported that InterComp tended to perform less well on small surfaces.

Table S11. Performance of Ab-Ligity and InterComp on heavy chain or light chain only paratopes, on the full set. Antibodies modelled using both VH and VL chains are labelled 'full antibody', while those modelled using a single VH or VL chain only are labelled 'single domain antibody'.

| | | | **Epitope Definition** | | | |
|---|---|---|---|---|---|---|
| **Modelling** | **Paratope regions** | | **Ab-Ligity** | | **InterComp** | |
| | | **Paratope Definition** | **Precision** | **Recall** | **Precision** | **Recall** |
| **Full antibody** | **Original** | **Ab-Ligity** | 0.95 | 0.85 | 0.83 | 0.92 |
| | | **InterComp** | 0.92 | 0.77 | 0.80 | 0.83 |
| | **VH paratope** | **Ab-Ligity** | 0.90 | 0.78 | 0.78 | 0.84 |
| | | **InterComp** | 0.75 | 0.80 | 0.65 | 0.85 |
| | **VL paratope** | **Ab-Ligity** | 0.64 | 0.90 | 0.54 | 0.94 |
| | | **InterComp** | 0.16 | 0.93 | 0.13 | 0.95 |
| **Single domain antibody** | **VH paratope** | **Ab-Ligity** | 0.88 | 0.78 | 0.76 | 0.84 |
| | | **InterComp** | 0.74 | 0.82 | 0.63 | 0.88 |
| | **VL paratope** | **Ab-Ligity** | 0.67 | 0.89 | 0.57 | 0.93 |
| | | **InterComp** | 0.17 | 0.94 | 0.14 | 0.95 |

### References

Kovaltsuk, A. *et al.* (2018) Observed antibody space: A resource for data mining next-generation sequencing of antibody repertoires. *The Journal of Immunology*, **201**(8), 2502-9.

Leem, J. *et al.* (2016) ABodyBuilder: Automated antibody structure prediction with data–driven accuracy estimation. *mAbs*, **8**(7), 1259-1268.

Liberis, E. *et al.* (2018) Parapred: antibody paratope prediction using convolutional and recurrent neural networks. *Bioinformatics*, **34**(17), 2944–2950.

Mirabello, C. and Wallner, B. (2018) Topology independent structural matching discovers novel tem-plates for protein interfaces. *Bioinformatics*, **34**(17), i787–i794.
